# NADPH diaphorase neuronal dystrophy in gracile nucleus, cuneatus nucleus and spinal trigeminal nucleus in aged rat

**DOI:** 10.1101/2019.12.21.885988

**Authors:** Wei Hou, Yunge Jia, Yinhua Li, Zichun Wei, Xiaoxin Wen, Chenxu Rao, Ximeng Xu, Fuhong Li, Xianhui Wu, Haoran Sun, Hang Li, Yizhe Huang, Jingjing Sun, Gege Shu, Xinghang Wang, Tianyi Zhang, Geming Shi, Anchen Guo, Shengfei Xu, Guanghui Du, Huibing Tan

## Abstract

NADPH-diaphorase (N-d) activity is commonly used to identify NOS-ergic neurons. In our previous study, N-d positive neuritic dystrophy and spheroid termed aging-related N-d Body is discovered in the lumbosacral spinal cord in the normal aging rats. Histological studies also reveal that N-d positive neurodegenerative changes occur in the gracile nucleus. We re-examined N-d activity in gracile nucleus in aged rat. We found N-d positive neuritic dystrophy and spheroid also occurred in the cuneatus nucleus and spinal trigeminal nucleus. Besides regular coronal section, longitudinal oriented dystrophic neurites were detected in the sagittal and horizontal section in gracile nucleus and dorsal column. We fziurther examined the medullary oblongata with regular classical histology including Golgi staining, immunocytochemistry of NOS and phosphorylated tau protein, neuronal tracing method with wheat germ agglutinin conjugated alexa-fluor-488 through sciatic nerve, and spinal cord transection at thoracic level. Most of N-d positive neuritic dystrophy and spheroid did not showed colocalization with NOS or phosphorylated tau protein. Neuronal tracing and spinal cord transection revealed that N-d dystrophic neurites in gracile nucleus originated from terminal of sensory projection from spinal cord and peripheral somatic input. The results suggested that aging-related N-d dystrophy in the gracile nucleus was unique morphological feature. In conclusion, it was postulated that the N-d dystrophy as a morphological marker of aging degenerative damage in normal aged organisms.

## Introduction

The dorsal column nuclei contain the neuronal terminals ascending from the first-order afferent neurons in the dorsal root ganglia (DRG) and second-order afferent neurons which are postsynaptic neurons relayed from the first-order afferent neurons. The terminals of the first-order afferent neurons are also termed as the distal fibers of the primary sensory neurons in which could innervate both somatic trunk and extremities [1–3] as well as pelvic organs [4–6]. A range of diverse functions of the gracile nuclei are studied by neurobioactivative examinations [7–11]. Nicotinamide adenine dinucleotide phosphate diaphorase (N-d) positivity is detected in the gracile nuclei[12]. Previous studies demonstrate that N-d positivity are identified to nitric oxide synthase (NOS) within the central and peripheral nervous system [13–15]. N-d positive neurons and axons projecting via gracile fascicle to gracile nucleus have been identified in the sensory pathway in the spinal cord [16, 17], DRG[18] and gracile nuclei [19]. N-d staining is considered to label NOS or non-NOS neuronal components to demonstrate neurondegeneration or neurodegenerative diseases [20–26]. However, morphologically heterogeneous N-d reactivity also indicates that N-d-positive neurons may not always co-localize with NOS-positive staining [21, 27–29]. Neurodegenerative changes occur in N-d positive neuronal components in aging rats [30, 31]. Axonal dystrophy in normal aging is substantially an aberrant process located in the terminal parts of the axon in the gracile nuclei [32, 33]. Gracile axonal dystrophy (GAD) are aging related and non-specific lesions that occur in a wide variety of human and animal diseases [34–37]. Preference of neuroanatomy correlation of GAD is postulated in the primary sensory terminals of lumbosacral rather than cervical sensory neurons[38]. Gracile neurons present hyperexcitability to somatic noxious pain stimuli in aged rats [39] and receive visceral sensory inputs of pelvic organs. Neurodegnerative spheroids in gracile nuclei of animal models displayed some unique changes [40–42].

In our unpublished study in aged rats, the aging related NADPH-d neurodegeneration was not only detected in the gracile nucleus, which is similarly derived from the aging alterations in the lumbosacral spinal cord [21, 43], but also detected the cuneatus nuclei and spinal trigeminal nucleus. The inputs of the primary sensory neurons of DRG could form axon bifurcation in the spinal cord. The bifurcating collaterals can terminal in the corresponding spinal segments and simultaneously ascend to dorsal column respectively[38, 44]. Beside somatic sensory inputs, dorsal column nuclei also receive ascending visceral sensory information [45]. The spheroidal N-d neurodegenerations in the lumbosacarl spinal cord and gracile nucleus was postulated for dying back in aged rats [46–48].

Our previous finding indicated that aging-related N-d positive neurodegeneration has been found in the lumbosacral spinal cord[21]. Aging changes of neuronal soma and proximal dendrites are considered irregular swelling and “lumpiness” of cell bodies and initial portions of the dendrites[49]. The distal terminals of N-d positive abnormalities of the gracile nuclei are still lack of detailed demonstration in aging rats. Recently, our preliminary experimental results revealed that aging related N-d abnormobilities in gracile nuclei and lumbosacral spinal cord were non-somatic neurodegenerative structures. We hypothesized that the N-d neurodegeneration in gracile nuclei and lumbosacral spinal cord were distal terminals of primary sensory neurons from the DRG or bifurcating components in lumbosacral spinal segments. By horizontal or sagittal oriented sectioning, N-d labeling dystrophic neuritis were clarified as a swelling terminal in gracile nuclei. In this study, we will provide evidence that N-d positive neurodegeneration in the gracile nuclei are different from other neurodegenerative features.

## Materials and Methods

### Animals and tissue preparation

Young and aged male Sprague Dawley rats from 2, 4, 5, 8, 12, 18, 24 and 30 months were used for the experiments. Animals were raised in plastic cages with standard commercialized soft bedding and with adequate food and water at room temperature (22±1°C) and with a 12-h light and dark cycle (dark cycle 8:00 PM–8:00AM). Water and food were provided *ad libitum*. The animal experiment protocol was approved by the Animal Care and Use Committee in Jinzhou Medical University. Animals were deeply anaesthetized with pentobarbital 200 mg/kg and then intracardiacally perfused with normal saline followed by paraformaldehyde 4% in 0.1 M phosphate buffered saline (PBS, pH 7.4). After perfusion, the medullas and spinal cords cervical to coccygeal and were removed and postfixed in 4% paraformaldehyde overnight at 4 °C. The tissues were next transferred into 25% sucrose in 4% paraformaldehyde PBS for cryoprotection. Medulla and spinal cord segments were sectioned either coronally, horizontal or sagittal of 40 μm thick section using a cryostat (Leica, German).

### N-d staining

The medulla and spinal cord sections were processed for N-d activity according to the procedure of Tan 2006[21]. Briefly, sections were incubated in 0.1 M PBS pH 7.4, containing 0.3% Triton X-100, 0.5 mg/ml nitroblue tetrazolium (NBT, Sigma, Shanghai, China) and 1.0 mg/ml β-NADPH (Sigma, Shanghai, China) at 37°C for 1-3 h. Following the reaction, the sections were rinsed in PBS, air-dried and coverslipped with Neutral Balsa (China). In order to further check neuronal cells with nuclei, neutral red (see following method) and DAPI was also used for counterstaining following N-d staining.

### Classical histological staining

The Golgi staining is still used for the visualization of degenerating axons and dendrites, arborization and connections [50–52].The dorsal column nuclei are relay nuclei of central projecting terminals of the primary sensory neurons from DRG. Recently, the dendritic loss of neurodegeneration is indicated by Golgi staining[53]. The classical histological staining including Golgi method to demonstration basic property of the aging neurodegeneration in the dorsal column nuclei.

### Golgi staining

Experimental animals of 2-month- and 24-month-rats (n=5 respectively) were deeply anesthetized and sacrificed by bilateral thoracotomy without perfusion. Experimental procedure was performed by a modified protocol according to a commercial Golgi staining kit (FD Rapid GolgiStain™ Kit, FD NeuroTechnologies, Inc, China). Briefly, the animal brain and spinal cords were quickly removed and carefully handled to avoid damage or pressing of the tissue. Tissues were rinsed quickly in double distilled and immersed in the impregnation solutions (solution A, B and C) according to kit protocol. Sections was cut on cryostat at thickness of 160 µm. Each section was transferred and mounted on gelatin-coated microscope slides and processed with solution C, D and E. After staining procedure, sections dehydrate in graded ethanol. The sections were cleared in xylene and coversliped.

### Nissl stain and neutral red stain

For Nissl staining, the frozen sections were stained with 0.1% cresyl violet for 10-20 min. Some sections of N-d staining were counterstained with neutral red. After histological staining, sections were rinsed PBS, dehydrated in a graded alcohol series, cleared with xylene and mounted with neutral gum. For neutral red staining, the sections were incubated with 0.1 % solution at room temperature for 1-10 minutes depending on the desired intensity. After incubation, extra solutions were washed out with dH_2_O.

### Double staining for N-d histology combined with immunocytochemistry

After N-d staining in medullary oblongata or spinal cord sections, the sections were processed with immunocytochemistry. For the immunohistochemistry, a free-floating method were performed. Briefly, sections were incubated with 1% H_2_O_2_ in methyl alcohol for 30 min at room temperature, incubated with 5% normal horse serum for 30 min at room temperature, then incubated with primary antibodies for 48 h at 4 °C. The sections were collected in PBS in 24-well plates and processed for free-floating immunocytochemistry using primary polyclonal antibodies of phosphorylated Tau pSer214 (, rabbit, 1:1000 Sigma, USA) and NOS (Cayman Chemical,1:1000, USA). They were then treated for 2 h at room temperature with biotinylated secondary antibody (Vector Laboratories, VECTASAIN ABC Kit). Washing between steps was rinsed with PBS. Streptavidin-Biotin-Peroxidase labeling was visualized by incubating with a solution containing 0.001% 3.30-diaminobenzidine and 0.0003% H_2_O_2_. When a dark brown product formed, the reaction was terminated. Sections were washed, dehydrated with graded alcohol, then coverslipped. In each immunocytochemical testing, a few of sections were incubated without primary antibody, as a negative control. No specified staining was detected in the immunostaining control procedure.

### Neuronal tracing with WGA alexa-fluor-488

Rats were anesthetized with intraperitoneal injection of 3% sodium pentobarbital solution (30-50 mg/kg body weight) and the right lateral thigh was prepared for surgery. The tracer was dissolved or suspended in sterile saline. The right sciatic nerve was exposed and injected with 5 μl of WGA alexa-fluor-488 (Invitrogen, USA), using a 10-μl microsyringe. The needle of the microsyringe was allowed to remain in place for 5 minutes and slowly withdraw to prevent tracer leakage. For control injection of tracer, sterile saline (5μl) was injected into the age match rat, and the exposed sciatic nerve was left intact in the sham surgery group. Incisions were properly closed and the rats were housed and fed ad libitum under a 12-hour light/dark cycle. Three-five days after intra sciatic nerve tracer application, animals were anesthetized with pentobarbital 200mg/kg and transcardially perfused with 50-100 saline followed by 500 ml of 4% paraformaldehyde in 0.1 M PBS (pH 7.4).

### Spinal cord transection at mid thoracic spinal cord

T7 dorsal laminectomies were performed. Rats (2-month, n=5 and 18-month n= 6) were deeply anesthetized by isoflurane inhalation with isoflurane anesthesia vaporizer (RWD, China). Briefly, the dura was opened. To make transection lesions, the dorsal cord midline was identified and superficially incised with microscissors. Incisions were properly closed After the operation. rats were housed singly, kept warm at 26°C for 30 days, fed ad libitum with a 12-hour light/dark cycle, and given manual bladder evacuation 3 times daily for a period of 10 d and intramuscular ampicillin (25 mg twice per day) to prevent and treat urinary tract infections.

### Figure edition

The sections were observed and photo images were taken using fluorescence microscope (Olympus BX35, camera DP80, Japan). Figures were edited by photoshop and CorelDraw X6.

## Results

### Morphology, distribution and orientation pattern of the N-d dystrophy in the medullary oblongata

In young adult rat (2 and 4 month), several neurons and neuronal fibers were visualized by the N-d enzyme histology. The N-d reactivity was distributed in broad regions and nuclei, such as the gracile nucleus, conatus nucleus and spinal trigeminal nucleus (Figure 1). In this study, the major finding was focused on aging alteration of N-d staining in the three regions, which related to primary sensory pathway. There was also constitutively other regional N-d positivity such as reticular formation, dorsal nucleus of the vagus nerve and solitary tract nucleus and so on.

**Figure 1.**
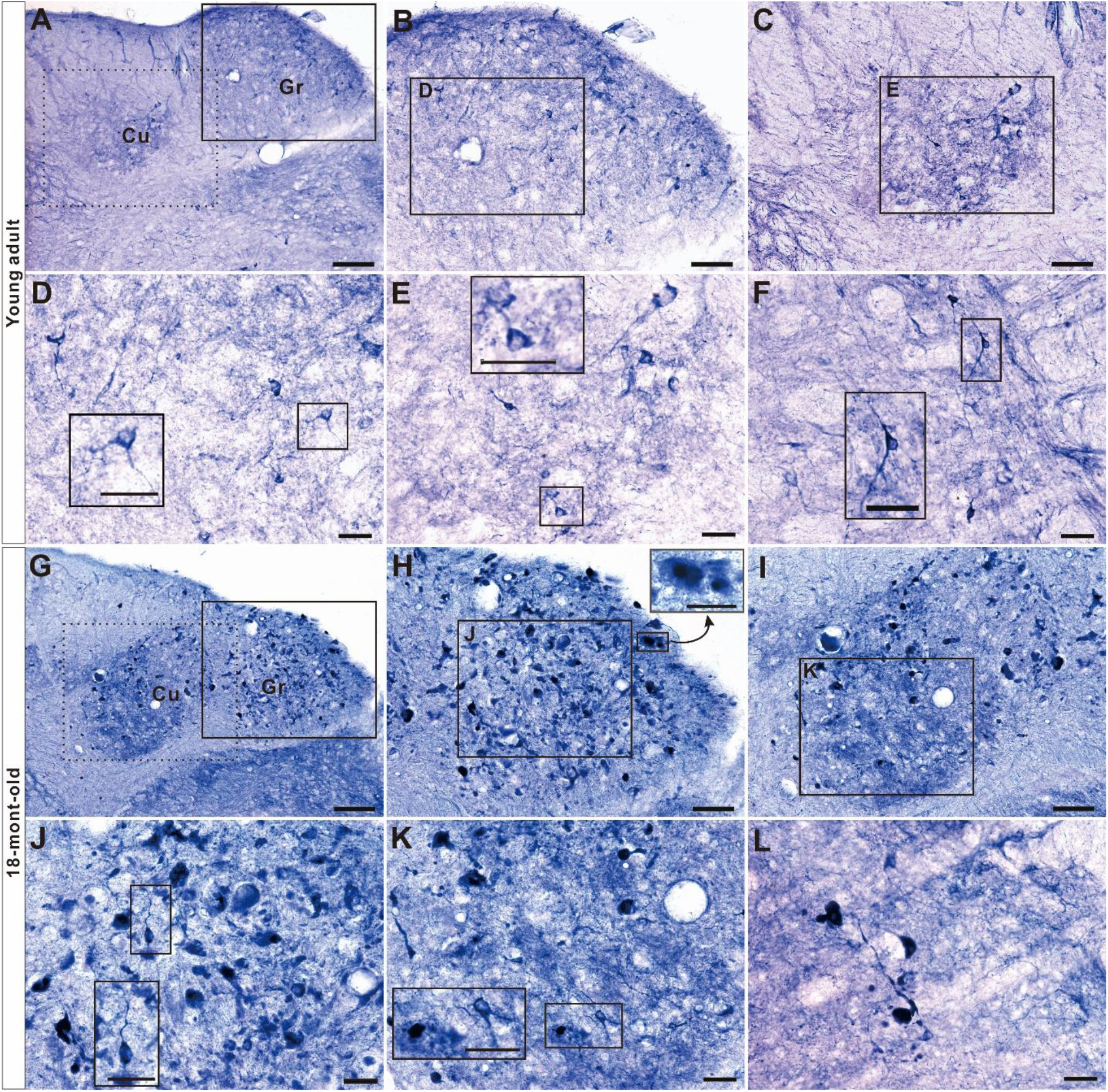
N-d positivity in the gracile nucleus, cuneatus nucleus and spinal trigeminal nucleus of young adult and aged rat. A: Gracile nucleus (Gr) and cuneatus nucleus (Cu) of young adult rat. B and C showed Gr and Cu of higher magnification from A respectively. D and E showed higher magnification from B and C receptively. The insets of D and C showed higher magnification of N-d positive neuronal soma. F showed spinal trigeminal nucleus of young adult rat. Inset showed neuronal soma. G: Aging-related N-d dystrophic spheroids detected in Gr and Cu of 18-mouth-old rat. H indicated higher magnification of G. Small rectangle showed higher magnification of two dystrophic spheroids with highly intensive stained core surround slightly stained halo. I indicated aging-related N-d dystrophic spheroids in cuneatus nucleus from G. J and K showed higher magnification of H and I respectively. Higher magnification in K showed both a neuronal soma and an aging-related N-d dystrophic spheroid also surround lightly stained halo. L showed aging-related N-d dystrophic spheroids in spinal trigeminal nucleus. Bar (A and G) = 100μm. Bar (B, C, H and I) = 50μm. Bar (D-F and J-L) =20μm.

Besides normal N-d activities of somatic component and neuronal fiber in the medullary oblongata, aging-related N-d alteration was detected in the gracile nucleus, cuneatus nucleus and spinal trigeminal nucleus, which revealed as non-somatic, dystrophic neurite and spheroid. N-d neurodystrophy in gracile nuclei occurred more than 6 months and increased with aging. In the coronal sections, the profile of the N-d neurodystrophy was spherical shaped or anomalous non-somatic structure. The diversity of the N-d morphological alteration could be visible in density with moderate or lightly stained, dark or dense staining, regular enlargement, enlarged or small dotted like spheroid, irregular or eccentric form moderate or lightly stained, dark or dense stained or cloudy like component (Figure 1). In the later adult-age (8 month), the dystrophic neurites firstly occurred in the medial part of the gracile nucleus and gradually occurred to lateral part with aging. After 18month, the distribution of the N-d neurodystrophic neurites were throughout the almost entire gracile nucleus from caudal to rostral levels and also from medial to lateral parts. spaciatempolly, the early dystrophic neurites were small sized-dot like abnormality and gradually dilatated to large spheroids with a certain diversity. The dystrophic neurites were detected earlier than that of the lumbosacral spinal cord (data not showed here). In the sagittal section from medial to lateral, it illustrated orientation of N-d dystrophic neurite. The neurogenerative spheroids increased in the rostral to gracile nucleus. Meanwhile, dilated neurites showed also increase in from caudal to rostral. In the Figure 2-1, the neurite alteration did not frequently occur in the dorsal column at most caudal end compared with that in the gracile nucleus. The medial section cross gracile nucleus still showed longitudinal neurites and variety of spheroids in both gracile nucleus in the same section of aged rat (Figure 2-2). In Figure 3, lateral section through cuneatus nucleus(rostral) and spinal trigeminal nucleus(caudal). In order to illustrate 3-dimensional information of the dystrophic spheroid and neurite. In the horizontal section, the profile of the N-d neurodystrophy was thread swelling neurite with dilated varicose neurodystrophy and clearly showed as nervous fiber terminals from the dorsal column fascicle of the spinal cord (Figure 4). The number of the N-d neurodystrophy was relative less than that of the gracile nuclei. Some of irregular eccentric N-d aberrant formation shaped like cloudy or massive plaque enormously occupied the parenchyma of the gracile nucleus. The lineated bead dystrophic spheroids or neurites still remained continuity, for example in Figure 2-1 c2 and d3.

**Figure 2-1.**
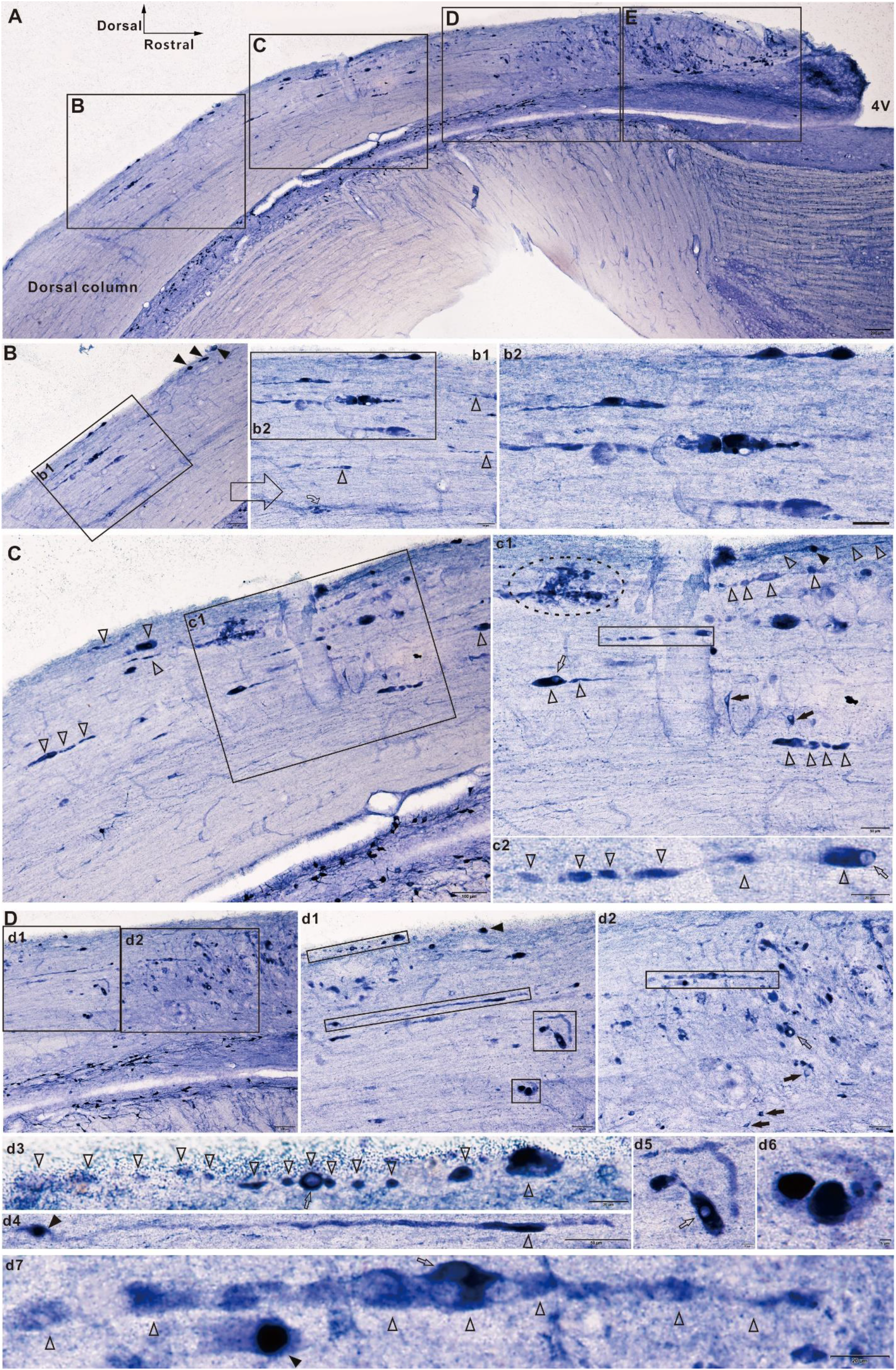
Orientation and diversity of N-d positive neurodegeneration in gracile nucleus and dorsal column in the sagittal section of aged rat. A showed sagittal oriented section from upper cervical segment of spinal cord to the gracile nucleus. 4V indicated the fourth ventricle. The four regions labeled B, C, D and E with rectangle outlines were presented in following, which were divided Figure 2 into 2 parts. E presented in Figure 2-2. B showed N-d positive dilated neurites in rectangle outline and swelling spheroids (Arrowhead) in dorsal column. N-d enlarged thread-like profiles showed with magnifying in b1 and b2. Curved arrow indicated cloudy-like N-d neurodegenerative profile. Open arrowhead dilated thread-like profiles. Several bead chain or dilated thread-like profiles showed in b2. C rostral to B showed increased dilated bead chain thread-like profiles. Linear arranged open arrowheads indicated dilated thread-like profile. c1 was magnification of C and showed some aligning linear dilated neurites. Dash line indicated a cluster of cloudy N-d alterative profile. Solid arrowhead with three open arrowheads indicated aligning linear dilated neurite with a spheroid in the middle. Open arrow indicated vacuolar spheroid extended thread-like profile (two open arrow heads). Arrow indicated neuronal cell. Many regular N-d positive fibers were detected. c2 was magnification of rectangle outline in c1. c2 showed bead chain neurite with vacuolar spheroid (open arrow). D clearly showed massive gracile nucleus in rostral part. Two rectangular out lines (d1 and d2) included several enlarged dilated neurites correlatedly occurred in dorsal column and gracile nucleus. d1 showed junction of dorsal column and gracile nucleus. Four rectangular out lines in d1 were marked for dilated neurites and magnified as d3, d4, d5 and d6. Arrow head indicated a spheroid. d3 showed disconnected spheroids (open arrowhead) in linear arrangement. A vacuolar spheroid was marked by open arrow. d4 was a dilated neurite with spheroid (arrow head). The staining intensity varied in the neurite. d5 showed two kind of neurites. A vacuolar profile was marked by open arrow. d6 showed two big strong stained spheroidal cores and two small light stained spheroids around halo-like positivity. d2 showed neurodegenerative neurites and spheroids in the gracile nucleus. d7 showed neuritic dilation with vacuolar spheroid (open arrow). Similar to d6, a spheroidal core around halo-like positivity (arrowhead) was also detected in d7 which was magnified from d2. Bar in A =200μm, bar in B,C and D = 100μm, bar in b1, b2, c1, d2 and d4 = 50μm, bar in c2, d3, and d7 −20μm, bar in d5 = 10μm and bar in d6 =5μm.

**Figure 2-2.**
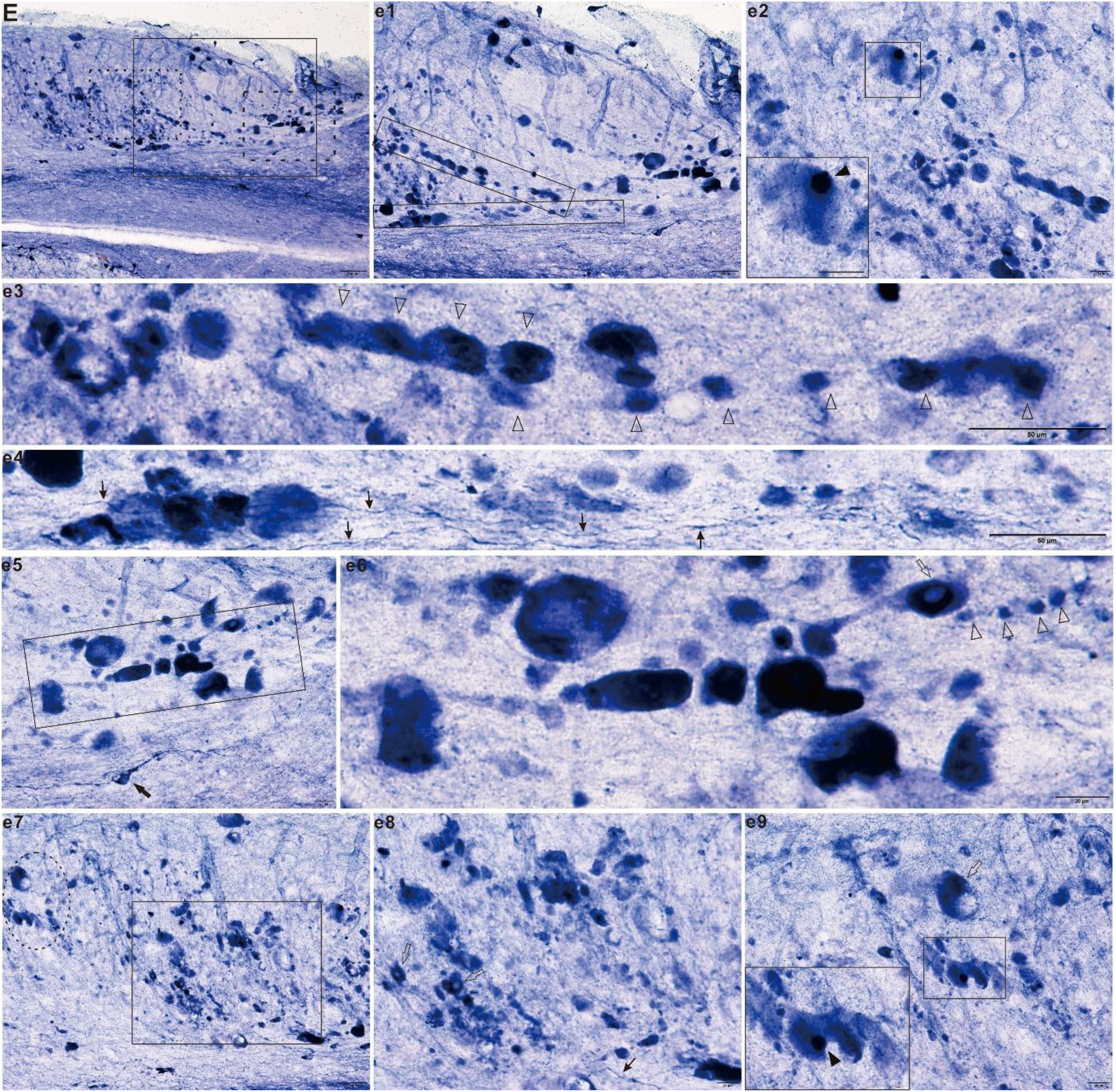
Orientation and diversity of N-d positive neurodegeneration in gracile nucleus and dorsal column in the sagittal section of aged rat (continued). All of the images were the same section showed in A from Figure 2-1. E was a magnification region of rectangular out labeled line in A. Some images overlapped and some magnified image extended out of boundary of E. They were all corelated of E. e1 was magnification of E. Several linear arranged swelling spheroids occurred in e1. Two of them showed in e3 and e4. e2 was magnification of dot rectangular out line in E. Squared out line and inset of e2 was a spheroidal core and halo. Two linear arrangements of neurodegenerative neurites (serial open arrow heads) showed in e3. Some regular neurites (thin arrow) occurred in e3. e5 was magnification of dashed rectangular outline. Arrow indicated a neuron with clear processes. e6 was further magnification from e5. Arrow indicated vacuolar spheroid. Open arrowhead indicated beat chain-like degenerative neurite. e7 showed many cloudy-like neurodegenerative profiles. e8 was magnification of e7. Open arrow indicated vacuolar spheroids. Thin arrow indicated regular fiber. e9 was magnification of dash line. Open arrow indicated vacuolar spheroids. Inset showed a spheroidal core and halo. Bar in E = 100μm, bar in e1, e3, e4 and e7= 50μm, bar in e2, e5, e6, e8 and e9 = 20μm, and bar in inset of e9 = 10μm.

**Figure 3.**
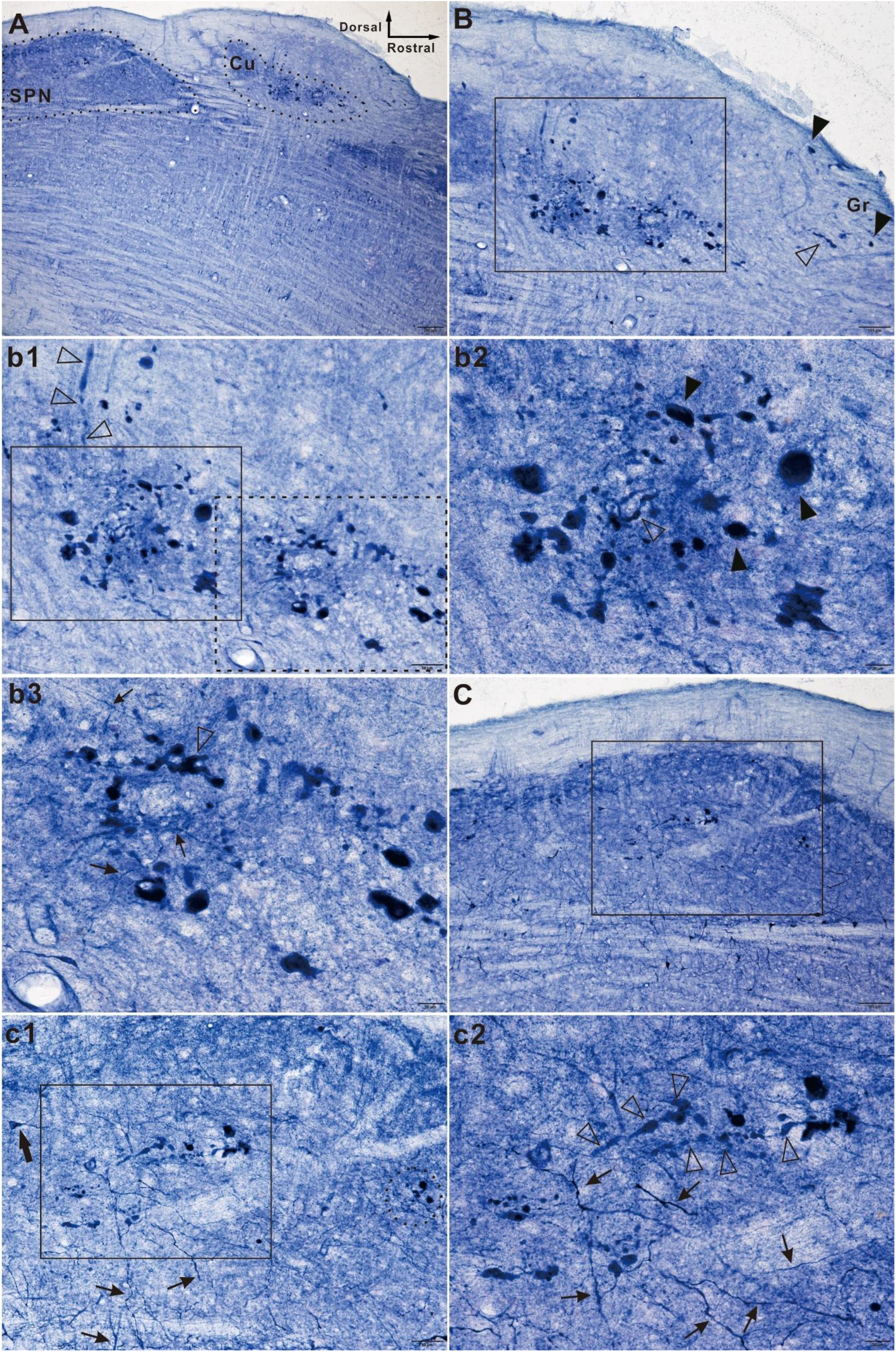
N-d positive neurodegeneration in cuneatus nucleus and spinal trigeminal nucleus in the sagittal section of aged rat. A showed sagittal section cross rostral cuneatus nucleus (Cu) and caudal spinal trigeminal nucleus (SPN) indicated with boundary of dot rectangular outline. B was magnification of Cu region from A. Some spheroids(arrowhead) and dystrophic neurite (open arrowhead) in dorsal rostral showed gracile nucleus (Gr). b1 was further magnification from B. Open arrow head indicated dilated neurites from dorsal to Cu. b2 showed further magnification from b1(rectangle). Arrowheads were example of spheroids. Similarly, open arrow head indicated twisted swelling neurites. b3 showed further magnification from b1(dash line). Thin arrow indicated regular neurite besides dystrophic neurite (open arrow head). C was magnification focus on SPN from A. Some normal regular neurites were detected in c1 in which also showed both rectangle and dot circle indicated dystrophic neurites. c2 was high magnification from c1. Bar in A =200μm, bar in B and C =100μm, bar in b1 and c1=50μm, b2 and bar in b3 and c2 −20μm.

**Figure 4.**
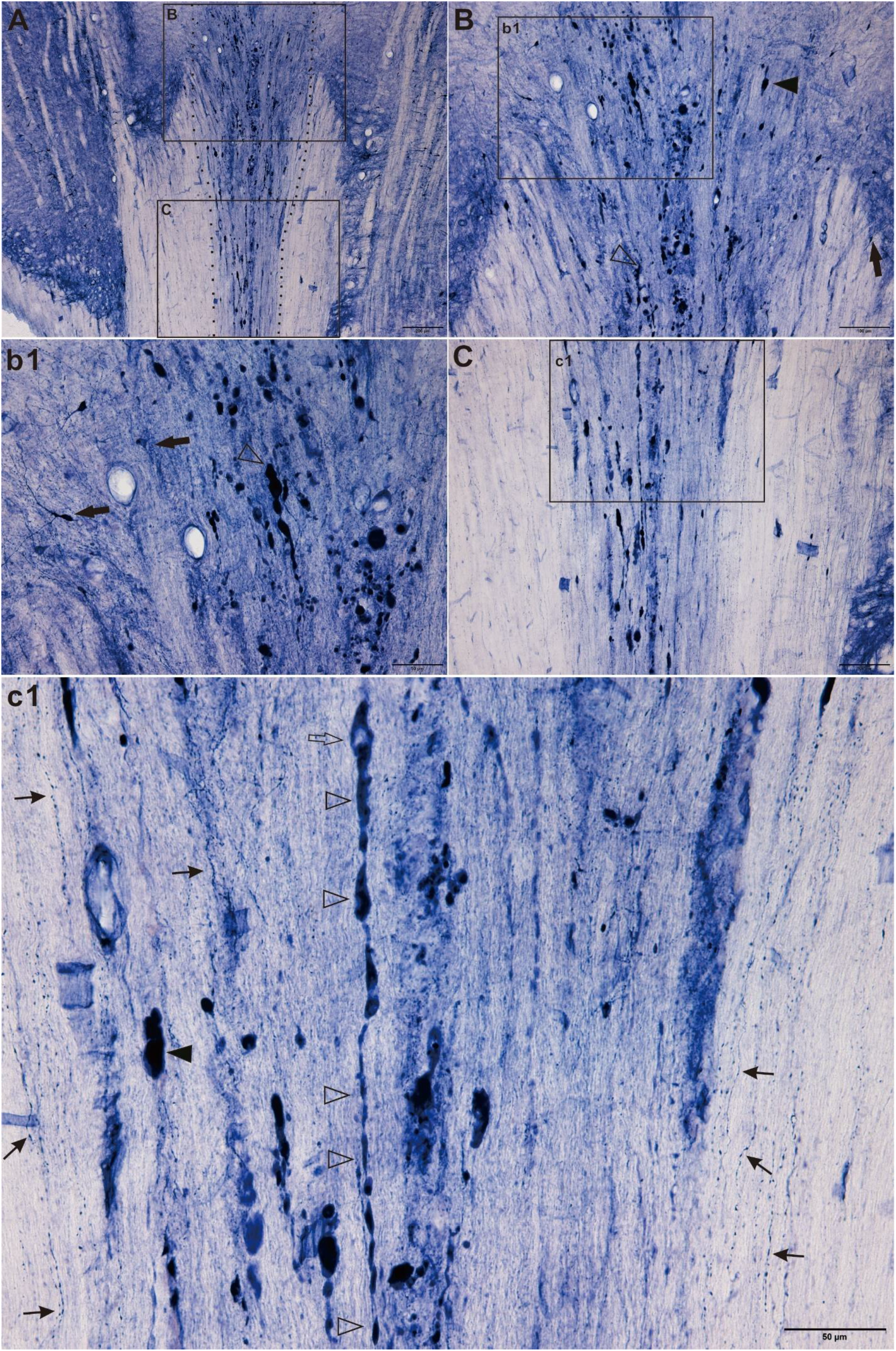
N-d histology in horizontal section for gracile nucleus in aged rat. A showed a part of gracile nucleus between dash line. Two rectangles presented in B and C. B showed rostral of the A in massive N-d dystrophic neurites. Example of spheroid(arrowhead), dystrophic neurites (open arrow head) and neuron(arrow) showed in B. b1 was magnification from B. C was magnification from A. c1 was magnification from C. A longitudinal swelling neurite indicated by linear open arrow head. Thin arrow indicated regular neurite. Arrowhead indicated example of spheroid. Bar in A = 200μm, bar in B and C =100μm and bar in b1 and c1 = 50μm.

The neurodystrophic structures could enlarged swelling of neuronal terminals. The constitutive neuronal tissue should be tested by traditional histology. Golgi staining showed decrease in neuronal fibers in the gracile nucleus in the aged rat (24-month-old). Meanwhile, spheroidal profiles of neuronal terminals were detected in the aged rats (Figure 5).

**Figure 5.**
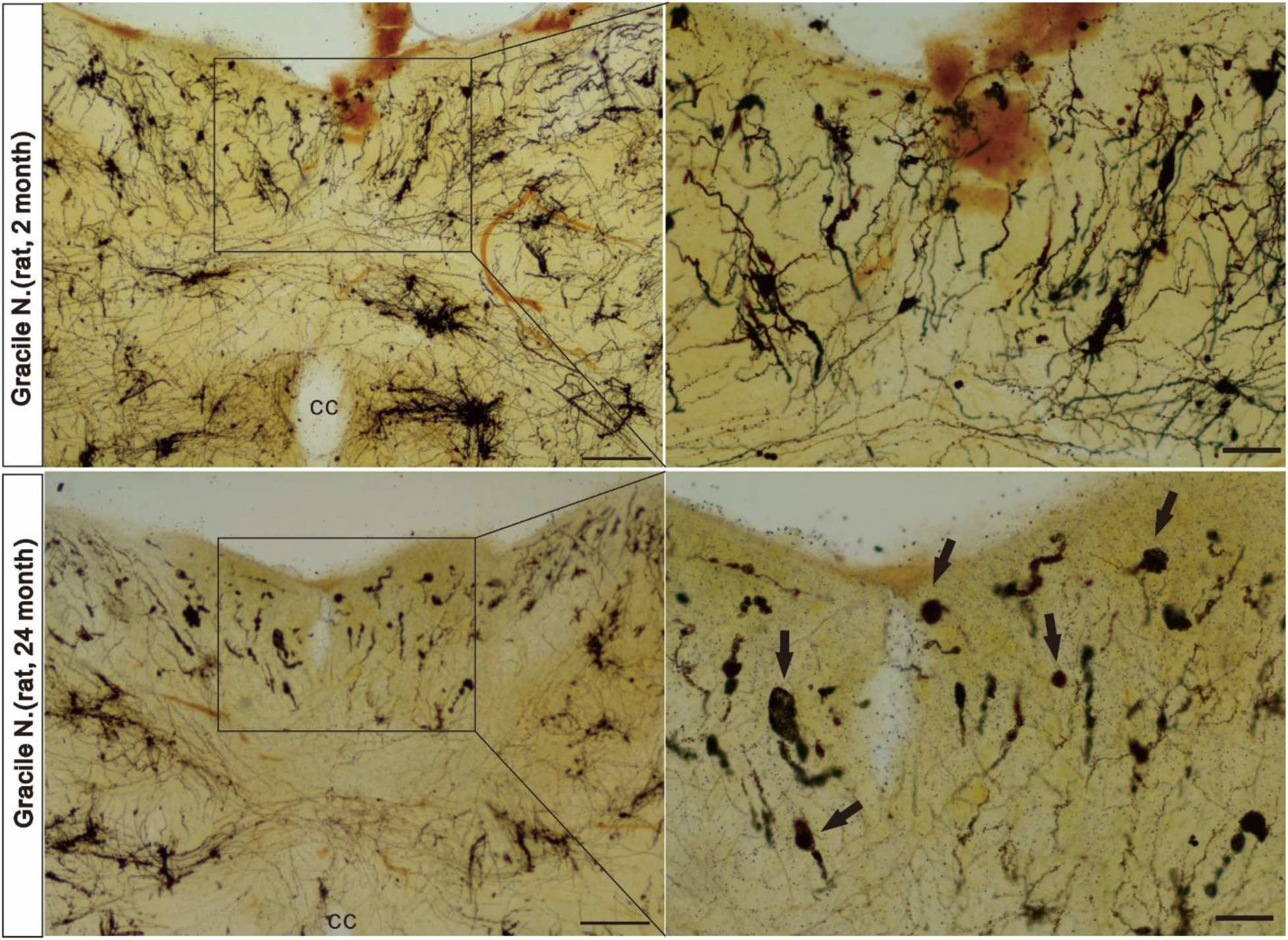
Morphological alteration of the gracile nucleus by Golgi staining. The decrease of neuronal fiber density was obvious and spheroidal profiles occurred in the aged rats compared with that of young rats. Bar for lower magnification=100μm, higher magnification=20μm

Different to Golgi staining, Nissl stain did not define any clear distinct dystrophic neurite and spheroid both in gracile nucleus (figure 6) in aged rat. Neutral red staining alone also did not clearly define the distinct dystrophic alteration, compared with counterstained after N-d histology in the aged rats (figure 6). Next, several sections of N-d dystrophic alteration were counterstained with DAPI. Figure 7-1 and Figure 7-2 showed that dystrophic neurite and spheroid in the gracile nucleus and cuneatus nucleus were not visualized as somatic structure with DAPI positive nuclei. While neuronal soma in the reticular formation showed DAPI positive nuclei in N-d positive soma in medullary oblongata of the same rat. The normal spatial for nuclei were usually no N-d reactive NBT formazan, which occupied by DAPI indicated nuclei.

**Figure 6.**
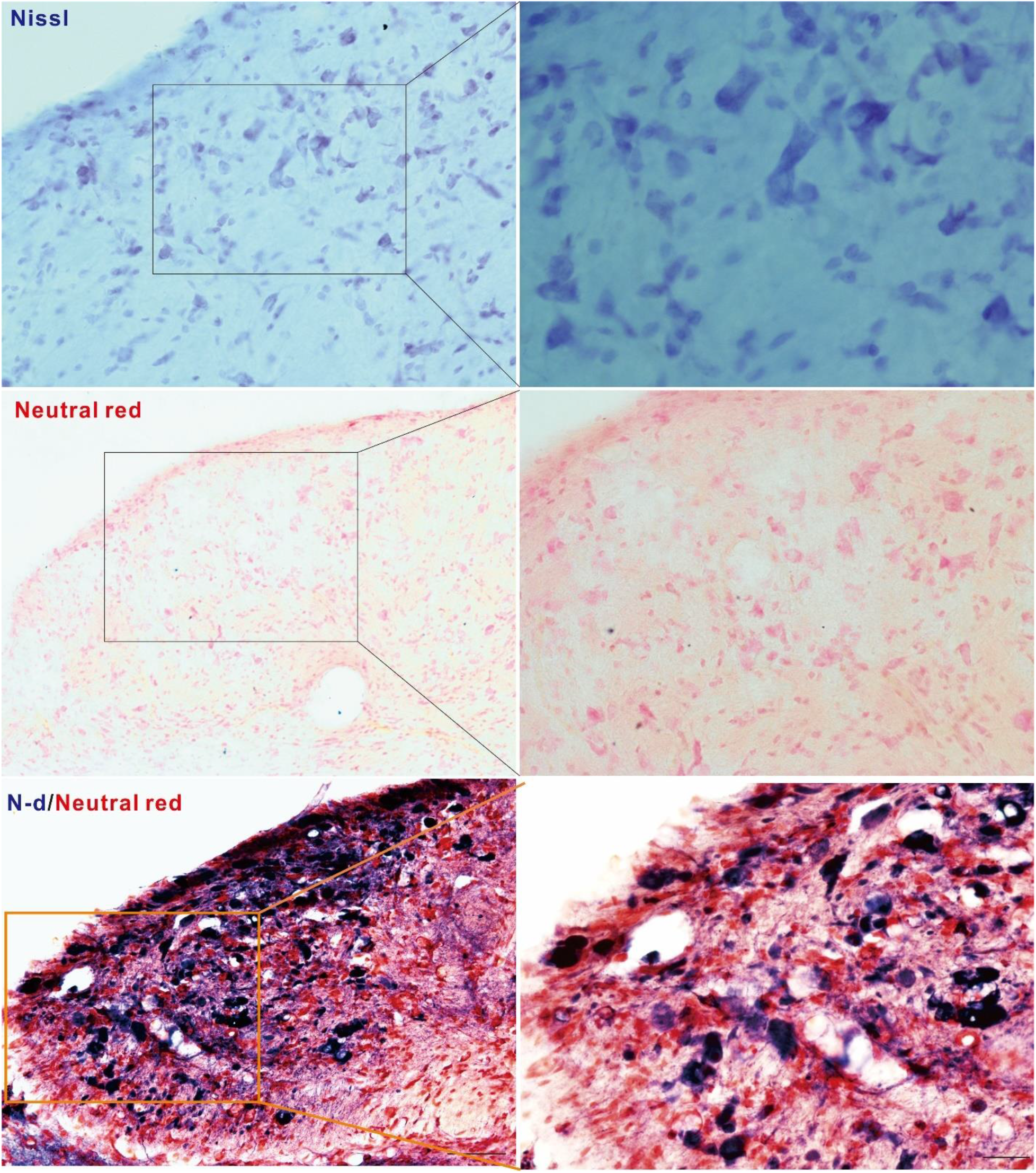
Nissl staining and neutral red staining as well as N-d enzyme histology with counterstained neutral red in the gracile nucleus. Left column is low magnification and right column is corresponding high magnification. N-d positive alteration for aging neurodegeneration scarcely stained by Nissl and neutral red. Bar for low magnification = 50μm, high magnification = 20μm.

**Figure 7-1.**
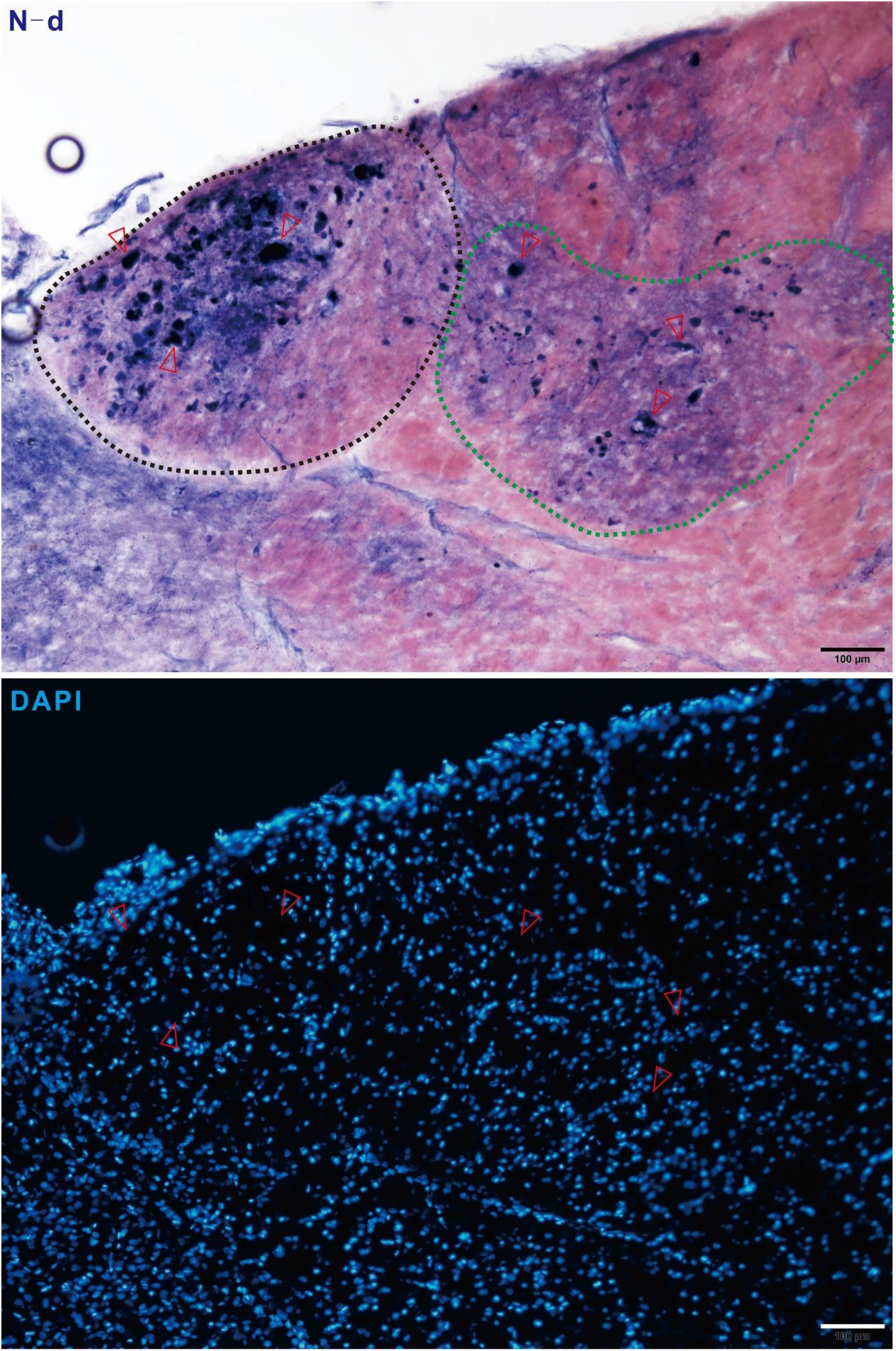
Another example of N-d staining of dorsal column nuclei in aged rat (24 months). A showed N-d positive spheroid in gracile nucleus (black dash line) and cuneatus nucleus (green dash line). Red opened arrows indicated typical large sized spheroids which also showed in B was lacking nucleus structures in the background of counterstained DAPI in same section. Bar = 100μm.

**Figure 7-2.**
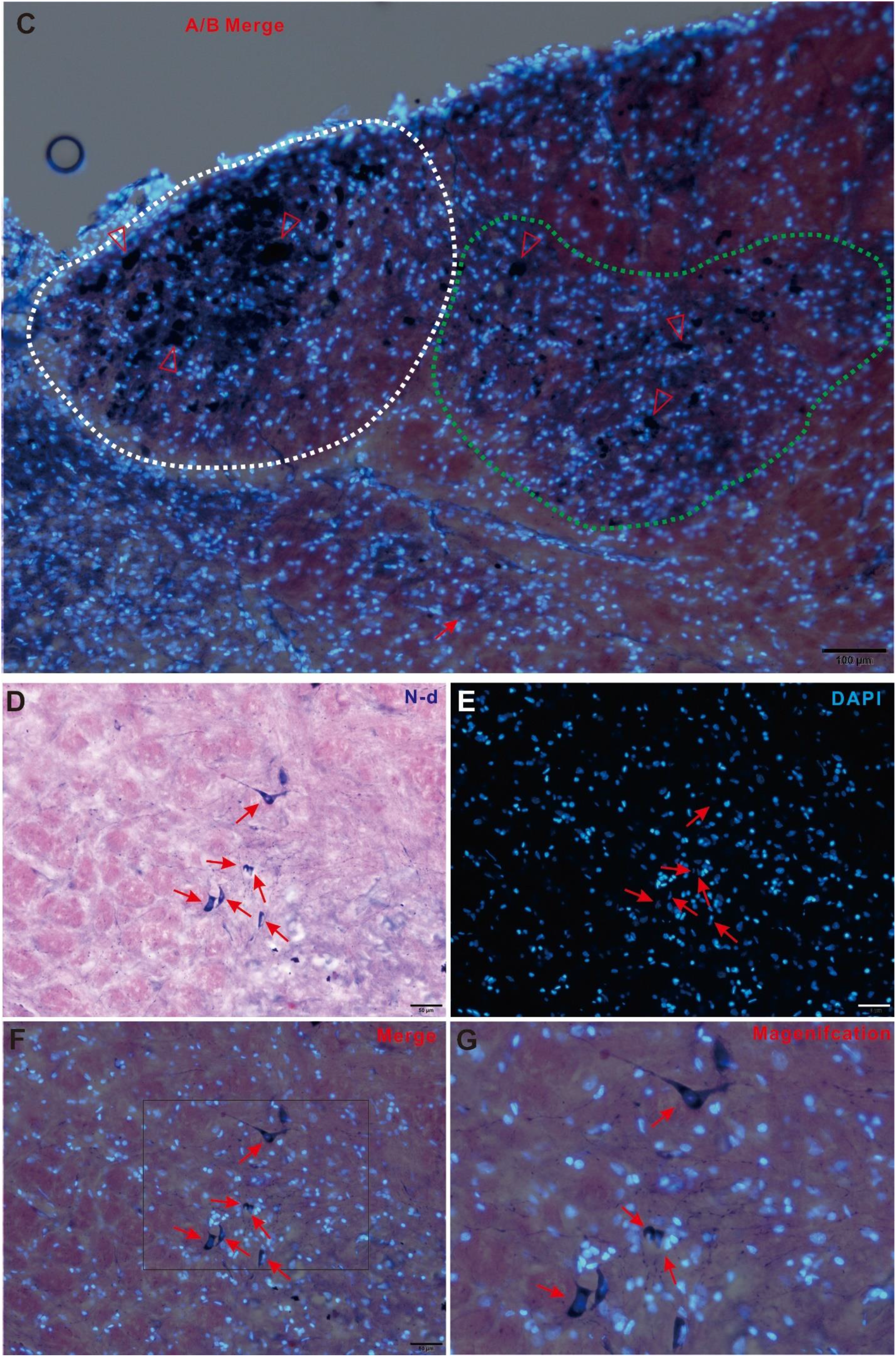
(Figure 7-1 continued) Non-somatic dystrophic spheroid in gracile nucleus and cuneatus nucleus and N-d positive neuron in reticular formation in the medullary oblongata of 24-month-old rat. C was merged image of A and B in Figure 7-1. Red open arrowhead indicated the same component of dystrophic neurite in the location in A and B in Figure 7-1. D-G was the similar staining pattern to A-C to show N-d positive soma in the reticular formation. Thin arrow indicated neuronal soma or nucleus of the same corresponding neuron. D showed N-d positive neurons (thin arrow). E showed counterstained with DAPI in the same location. F was merged images of D and E. G was magnification of E. Bar in C = 100μm and bar in D-G = 50 μm.

### NOS minor N-d dominant dystrophic profile

N-d positive neurons have been found to identical to NOS at various sites in the nervous system. In order to test the NOS associated with N-d positivity, we examined aging related N-d neurodegeration in gracile nuclei in aged rats. Basically, NOS positive neurons were visualized with N-d histology. We found that aging related N-d neurodegerative neuronal components may not always be NOS-positive in the gracile nucleus, cuneatus nucleus and spinal trigeminal nucleus. N-d enzyme histology of the spheroidal alterations partially co-localize NOS immunocytochemistry in aged rat (Figure 8).

**Figure 8.**
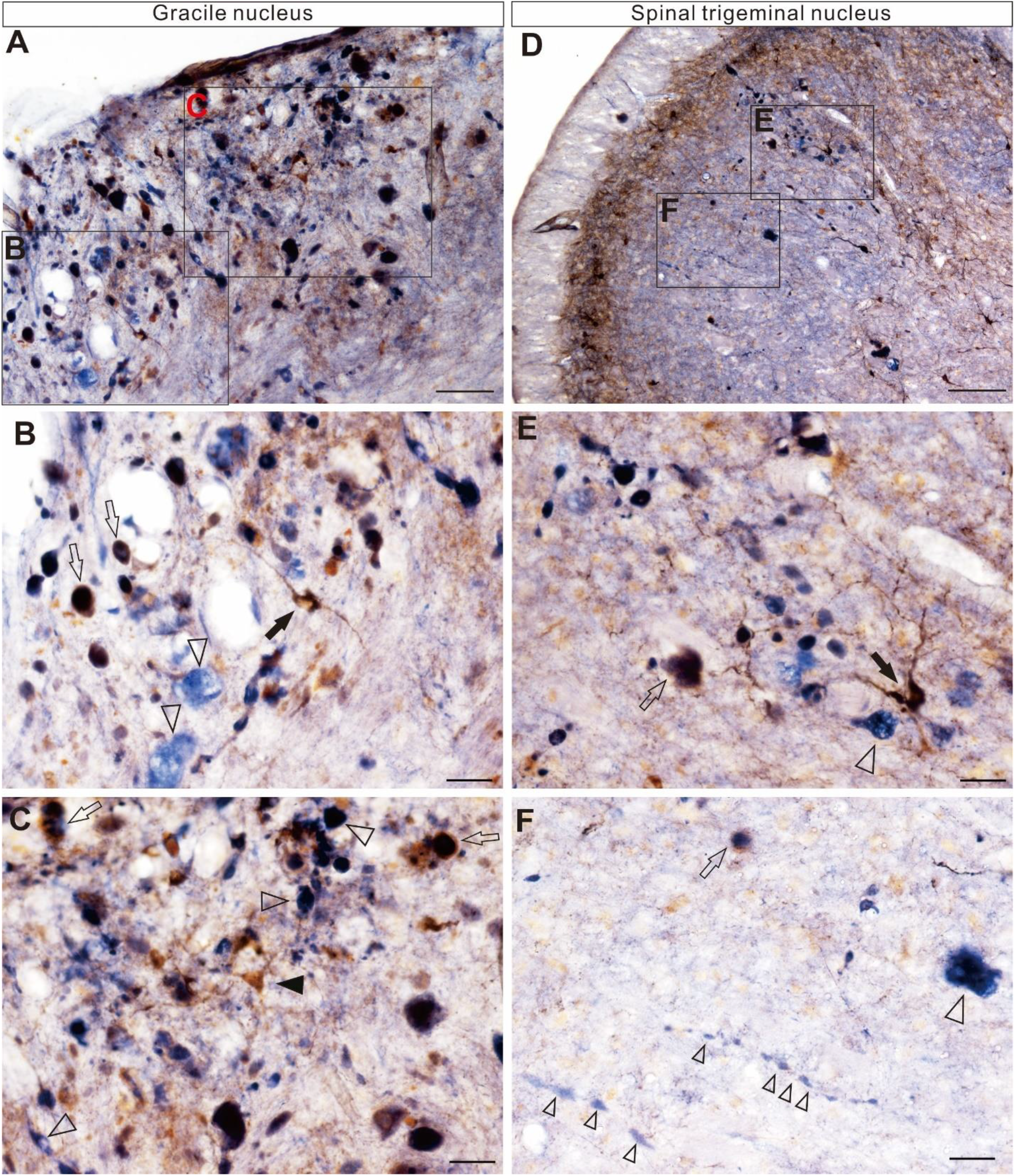
Double staining of N-d and NOS immunocytochemistry in the medullary oblongata of 36-month-old rat. A showed double staining for the gracile nucleus. B and C were higher magnification from A. Arrow indicated double stained neuron. Open arrow indicated double stained spheroids. Open arrowhead indicated single N-d staining. D showed double staining for the spinal trigeminal nucleus. E and F were high magnification from D. There were bead chain-like aberrant N-d positivity or enlarged varicosity-like structures (linear small open arrowheads). The size of many spheroids was much larger than that of the cells (B, C and E). Bar for A and D =50μm, and bar for B,C, E and F =20 μm.

### N-d dystrophy relevant other abnormalities in medulla oblongata

In order to demonstrate the distinguish the normal aging change to other neuropathology of neurodegenerative disease, we examined Tau immunoreactivity in the gracile nucleus in aged rat. We found that phosphorylated Tau positively occurred spheroid in 18-month-old rat (Figure 9). Tau spheroid or neurite can be detected in the spinal cord of both in non-demented controls and Alzheimer’ disease cases[54]. Following examination, spheroid of phosphorylated Tau occurred in gracile nucleus. Most of Tau spheroids did not co-localize with N-d spheroids besides small proportion of double staining. Less amount of Tau dystrophy was also detected in the sacral spinal cord.

**Figure 9.**
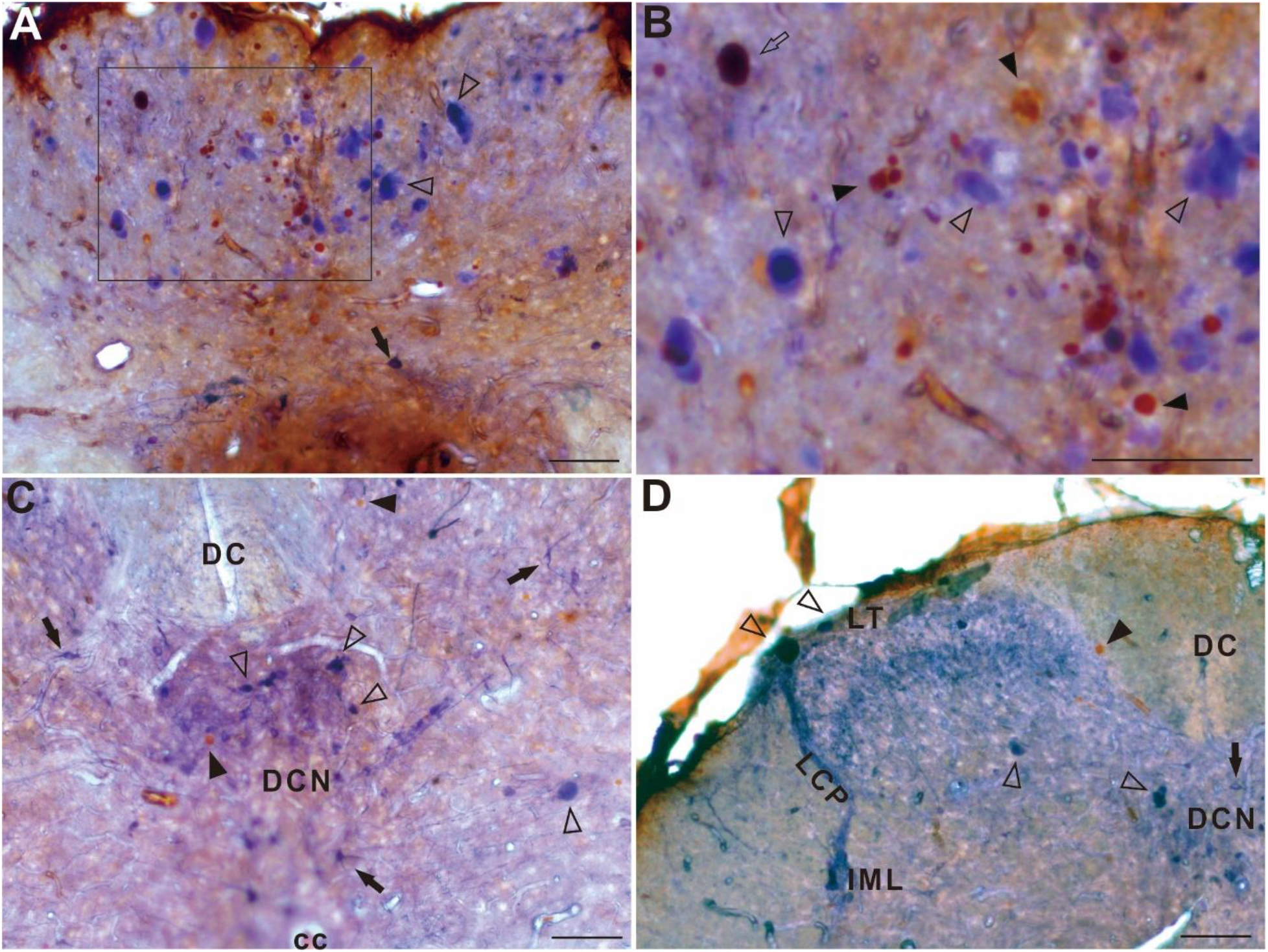
N-d histology combined with Tau immunoreactions in 18-month-old rat. A showed double staining of N-d histology and Tau immunocytochemistry. Arrow indicated N-d neuron. Open arrow indicated double stained spheroids. Open arrowhead indicated single N-d positive spheroids. C showed the double staining in the sacral spinal cord. DC: dorsal column; DCN: dorsal commissural nucleus; cc: central canal. D also showed double staining in the sacral spinal cord. LT: Lissauer's tract; LCP: lateral collateral pathway, IML: intermediolateral nucleus. sacral spinal cord. Bar for A-C= 100μm

Gracile nucleus is the termination of the primary sensory fibers from lumbosacral spinal nerve[2]. In order to tracing possibility of correlation between sensory input of lower extremities and the dystrophic neurites and spheroid in gracile nucleus. WGA alexa-fluor-488 was detected in the gracile nucleus of aged rat after injection of the tracer into sciatic nerve (Figure 10). There were some dystrophic neurite or spheroid showed the labeling of WGA alexa-fluor-488 from ipsilateral sciatic nerve, which indicated that dystrophic neurites were termination of primary ascending fibers from sciatic nerve. By mean of transection of spinal cord, ascending and descending of intraspinal and primary sensory projections can be studied in the rat spinal cord[56]. To further make a confirmation of ascending projection of N-d primary sensory fiber is corelated with dystrophic neurites in gracile nucleus. In our experiment, transection of spinal cord at the level of thoracic segment, the N-d positivity of dystrophic neurite most disappeared in gracile nucleus and the N-d positivity of dystrophic neurite and spheroid still showed no significant change in both cuneate nucleus and spinal trigeminal nucleus in aged rat (Figure 11). In young control, response to the transection of spinal cord at thoracic level showed degeneration response with decrease of majority N-d positivity of the processes of neurons in gracile nucleus (Figure 12). Meanwhile, cellular architecture of cuneate nucleus and trigeminal nucleus relatively maintained intact of N-d histology. Again, no aging-related N-d dystrophic spheroid and neurite were detected in the gracile nucleus, cuneate nucleus and trigeminal nucleus in young rats. The transection of the spinal cord at thoracic level in young rat did not result in the aging-related N-d alterations in the three nuclei, which was consistent with N-d histology in intact naïve young rats according to Figure 1.

**Figure 10.**
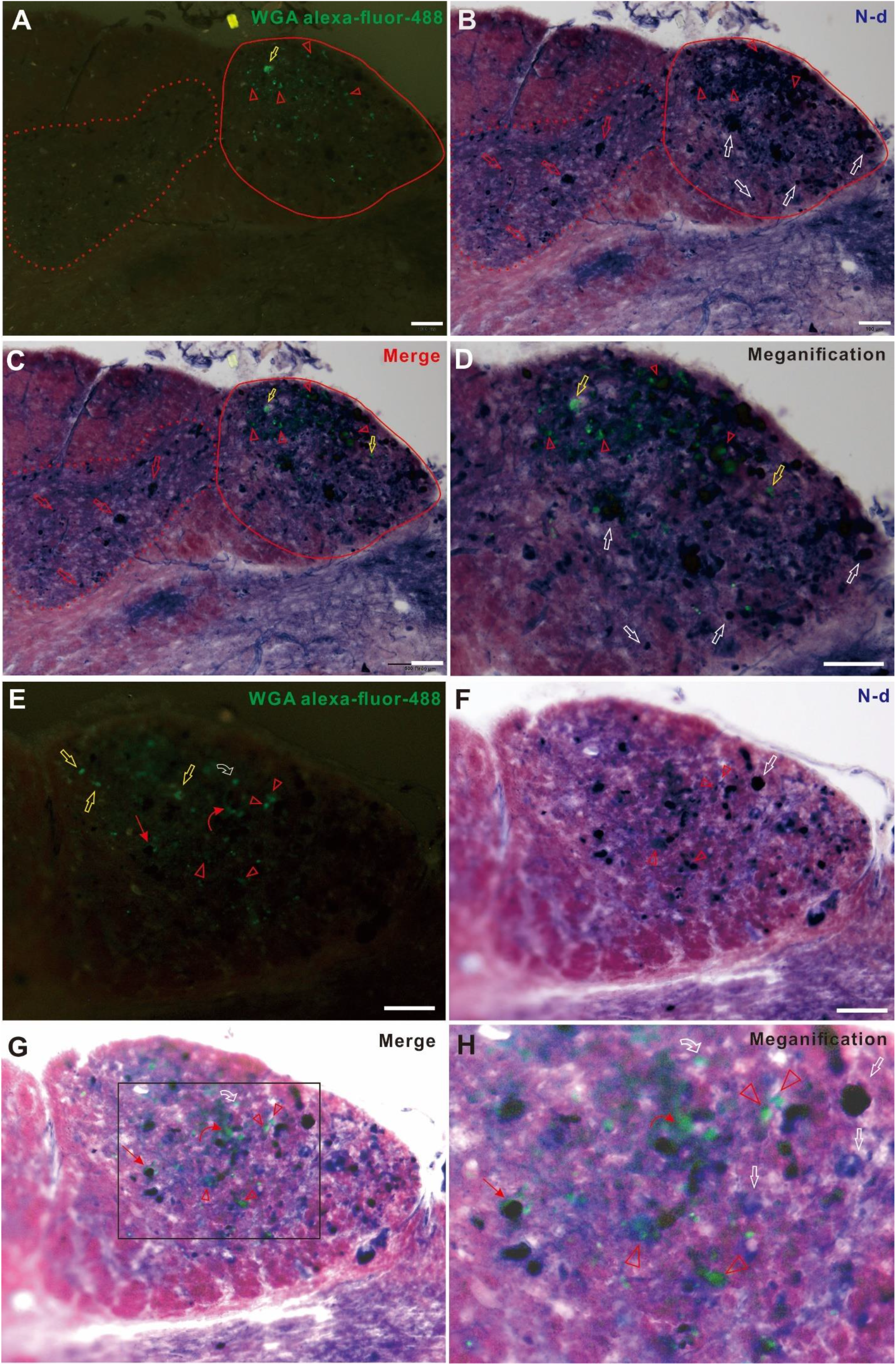
Positive terminals in gracile nucleus of 18-month-old rat labeled by injection of wheat germ agglutinin [55] alexa-fluor-488 into sciatic nerve. Open arrow heads pointed to the same locations in A, B, C and D. A showed positive labeling of WGA alexa-fluor-488 in gracile nucleus (red line) and no positive labeling in cuneatus nucleus (dot line). Yellow open arrow indicated example of N-d negative and WGA alexa-fluor-488 labeling spheroid. B showed N-d dystrophic spheroid in both gracile nucleus (red line) and cuneatus nucleus (dot line) as well as example of N-d dystrophic spheroid in gracile nucleus (white open arrows) and cuneatus nucleus (red open arrows). C showed merged A and B. D was magnification of C. E-H is another example for WGA alexa-fluor-488 labeling combined with N-d staining. E: Curved open arrow indicated WGA alexa-fluor-488 labeling spheroid surround N-d neurite. Thin arrow indicated WGA alexa-fluor-488 labeling fiber surround spheroid. Curved Thin arrow indicated WGA alexa-fluor-488 labeling neurite. F showed N-d staining spheroid (white arrow) and double staining spheroid (open arrow head) corresponding in G. G was merged image of E and F. H was magnification of G. Bar in A-D = 100μm and bar in E-H = 50μm.

**Figure 11.**
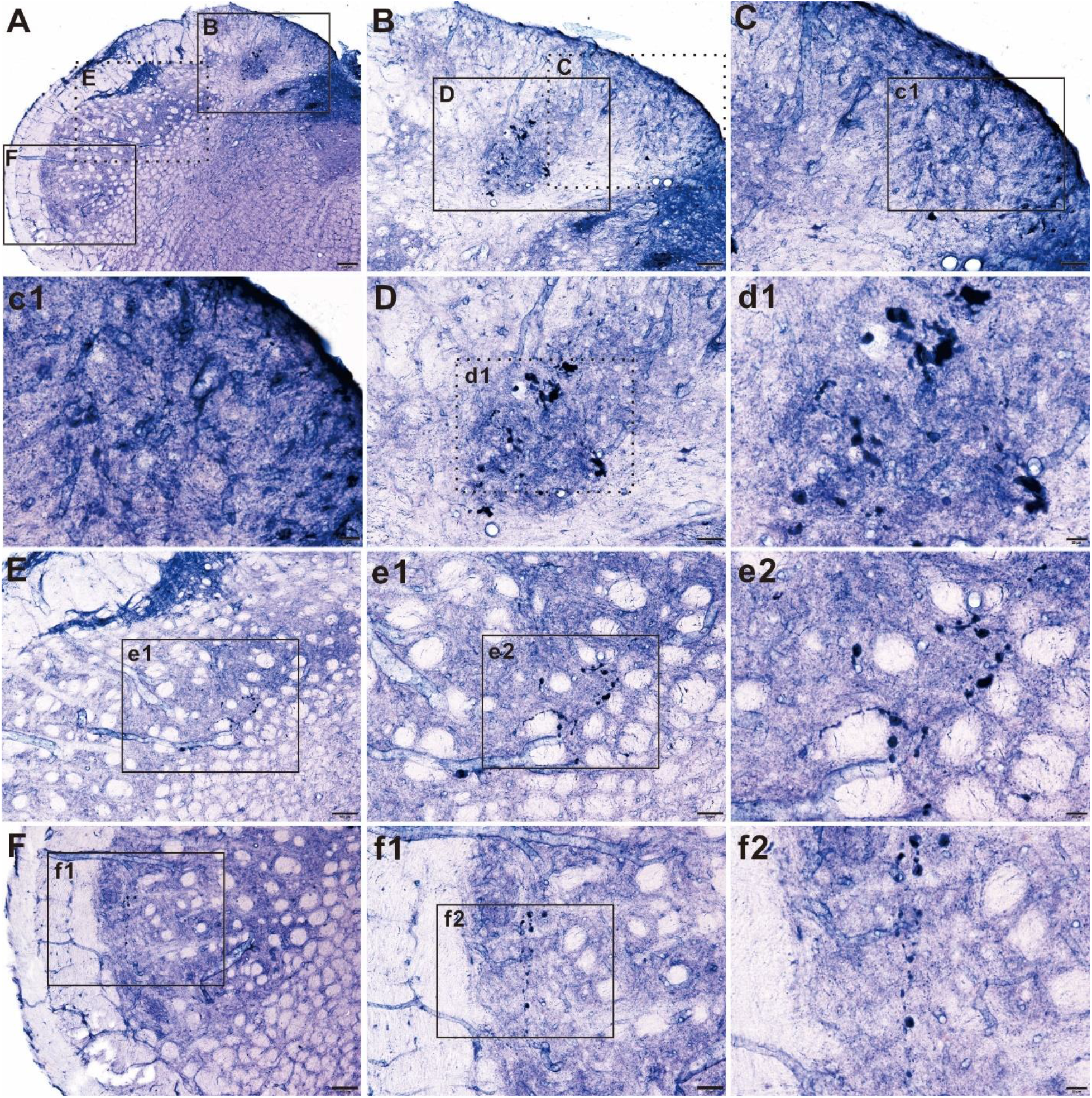
N-d reaction in medullary obligate in 18-month-old rat 1 months after transection of spinal cord at middle thoracic level. A: Rectangle B, E and F showed anatomy location for gracile nucleus, cuneatus nucleus and spinal trigeminal nucleus. B showed N-d dystrophic neurites and spheroids almost disappeared in gracile nucleus (most of dot rectangle C) and positive dystrophic stained in cuneatus nucleus (central region of rectangle D). C was magnification from B. c1 was magnification from C and no significant spheroid and dystrophic neurites was detected. Contrast to that in gracile nucleus, N-d dystrophic neurites and spheroids were still detected in cuneatus nucleus (D). d1 was further magnification from D. E showed N-d dystrophic neurites and spheroids in SPN from A. e1 and e2 were further magnification from E. Similar to E, F, f1 and f1 showed N-d dystrophic neurites and spheroids in SPN from A. Bar in A =200μm, bar in B, E and F= 100μm, bar in C, D, e1 and f1 = 50μm and bar in c1, d1, e2 and f2 =20μm.

**Figure 12.**
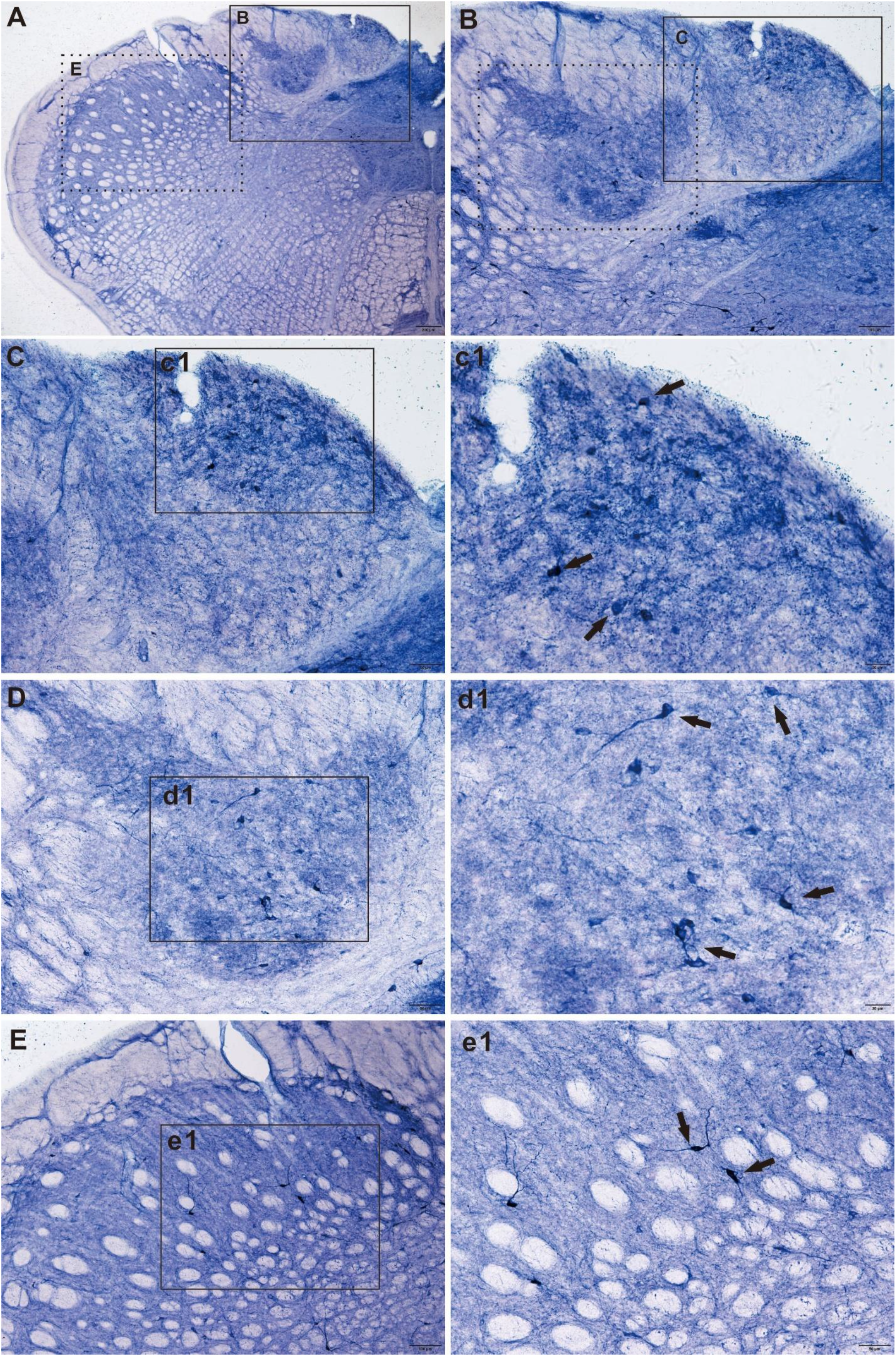
N-d reaction in medullary obligate in 2-month-old rat 1 months after transection of spinal cord at middle thoracic level. A: Rectangle B and E and F showed anatomy location for gracile nucleus, cuneatus nucleus and spinal trigeminal nucleus. B showed N-d histological staining in gracile nucleus (most of rectangle C) and positive stained in cuneatus nucleus (dot rectangle D). C was magnification from B. c1 was magnification from C and no significant spheroid and dystrophic neurites was detected. A few neurons were detected. The processes of neurons were not well stained as following neurons in cuneatus nucleus and spinal trigeminal nucleus. Similar to that in gracile nucleus, no N-d dystrophic neurites and spheroids were detected in cuneatus nucleus (D). d1 showed a few positive neurons(arrow). E showed regular normal N-d staining in SPN from A. e1 was further magnification from E. Bar in A =200μm, bar in B and E = 100μm, bar in C, D and e1 = 50μm and bar in c1 and d1 =20μm.

## Discussion

The present study has identified abnormalities of N-d positive dystrophic spheroid and neurite in the gracile nucleus, cuneate nucleus and spinal trigeminal nucleus in aged rat. We hypothesized that the dystrophic neuronal alteration was aging-related non-somatic components of distal ascending terminals of lumbosacral primary sensory neurons in aged rats. By means of horizontal and sagittal sectioning as well as analyzing of neuronal tracing experiment, firstly, we identified and characterized that N-d neurodegeneration of gracile nuclei occurred aged-dependent swelling neuronal terminals ascending from dorsal column of the primary sensory neuron from lumbosacral spinal cords. Secondly, we found that the N-d neurodegeneration were not completely co-localized with cNOS. Thirdly, we characterized the N-d neurodegeneration were not consistent with other common neurodegenerative pathology by immunocytochemistry of phosphorylated Tau protein. In further next experiment of neuronal tracing method, we illustrated that a part of dystrophic neurite originated from lower extremities. The result was reconfirmed with the degeneration method. To our best knowledge, we first demonstrate that N-d aging-related neurodegeneration also occurred in cuneatus nucleus and spinal trigeminal nucleus. Neurodystrophic alteration not only occurred in N-d positivity but also partially corelated with Tau neuropathology. We thought that all the neurodegenerative structures were N-d positive primary sensory central projecting terminals. Ascending fibers in dorsal column could be bifurcation from segmental primary sensory neurons[38, 57].

Both the gracile nuclei and cuneatus nuclei are belong to the dorsal column nucleus which locates in the dorsal and caudal of the medullar oblongata. Both nuclei contain second-order neurons of the dorsal column-medial lemniscus pathway, which carries fine touch and proprioceptive information from the body to the brain. The dorsal column nuclei receive inputs from sensory neurons of the DRG and send axons that synapse into the thalamus. The gracile nucleus not only participate in the sensation of fine touch and proprioception of the lower body (legs and trunk), but also carry the visceral sensory information [5, 45, 58].

### Relevant histologic and neuroanatomical features of the dystrophic alteration

N-d histochemistry reveals a unique population of neurons[59]. N-d neurons are distributed of in medulla oblongata in human [60, 61], frog[62] and rat[63, 64]. There are many neurotransmitters distributed in both sensory neurons in DRG and input perceive region in gracile nuclei[65–67]. N-d-positive neurons in DRG of the rat innervate viscera and visceral ganglia[68]. Gracile nucleus receives input from central projection of sciatic nerve[69]. Sciatic nerve injury directly causes responses of N-d expression in the gracile nucleus to in young and aged rats[70] as well as mid aged rat[71].The transganglionic degenerative changes such as complement activation are also induced in the gracile nucleus[72]. Ma et al report that strongly stained large and medium-sized cells, dot-like multipolar in shape, and large spheroids by N-d staining are detected in the gracile nucleus of aged rats[70]. In their previous study, a similar structures are termed as N-d-containing neuronal cell bodies[30]. The other types of swollen dystrophic elements are also detected with neuronal immunocythochemistry study[73, 74]. In our present study, the strongly stained large and medium-sized spheroids were considered as one kind of neurodegenerated dystrophic neurites, which also were detected in the cuneatus nuclei and the rat trigeminal nucleus. While, aging related N-d neurodegeneration has been revealed in the lumbosacral spinal cord of aged rats[21]. The N-d neuritic dystrophy in our present finding is a preferential degeneration of distal regions of longer fibers in the medulla oblongata, which is similarity to “dying back”[46, 47, 75]. For the primary sensory fibers, the longer ascending projection from the hindlimbs was presumed more vulnerable to neurodegeneration than the shorter one of upper limbs. Of course, the pathway of the sensation related trigeminal nerve is much shorter than that of That is why the more the dystrophic neurite and spheroid occurred in the gracile nucleus. The N-d dystrophy was detected only in small portion of spinal trigeminal nucleus in aged rat. In general, spinal trigeminal nuclei innervate the viscerotopic representation of the upper alimentary tract in the sensory ganglia of the IXth and Xth cranial nerves and in the subnuclei of the solitary and spinal trigeminal tracts[76]. It still needs to study the correlation between dorsal column nuclei with Spinal trigeminal nucleus[77–79]. Afferent inputs of sensory sciatic nerve can be detected in the dorsal lateral medulla in the rat, in which, neuronal tracing reveals a partial overlapping region[80].

### Axon dilation

Micromechanical action can cause reversible varicose in the axons of central neurons[81]. And neuropathology condition can also result in the dynamic neuritic change[82]. This phenomenon of physiology may also evident for varicosity of N-d positivity. Aging-related bead-like swellings and spheroid as well as dystrophic neurite could result in accumulation of normal proteins or aggregation of misfolding proteins. Tau pathology may occur in not only Alzheimer’s disease, but also Parkinson disease, prion disease as well as some cases of normal human[83, 84]. We noticed that tauopathy was also detected in gracile nucleus[85], and spinal cord[52, 54]. In our present investigation,

Some individual dilated axon spheroids seemed disconnected but it may still lineate alignment aligned as continuity. We presumed that molecule of the bioactive N-d dynamically moved in the axon and N-d aggregated into separated sites that formed spheroids. But these spheroids may still connect by axoplasm with ensheathed membrane. However, to a certain extent, the neurite or axon dilation finally membrane broken caused membrane leakage. We supposed that N-d left regular functional position in the neuronal plasma during the axon dilation. The intra cellular N-d spread to extra cellular space. N-d dystrophy of spheroid and neurite was disturbance for the abnormal N-d distribution. The shape of N-d dystrophy was variable such as dilated neurite, spheroid, dense core formation, vacuole formation, cloudy formation and spongiform. Widespread distribution of N-d dystrophy in the dorsal column nuclei may implicated to aging-related limb dysfunction, especially lower limb impaired movement. For severely dystrophic axons, some type of morphological profile is similar to spongiform plaque which may prion protein[86–88]

### Axon leakage

For diversity of intensive staining, at least three subclass N-d reactivity revealed: one lower intensive staining, another higher intensive dense staining and mixed staining. Some of the dystrophic formation had clear boundary and some formed cloudy blunt boundary. “Axonal leakage” is a novel type of axonopathy which usually accompanied with the extensive swollen axons and varicosities, and was associated with the origin and development of neurodegenerative disease [89–91]. In our present study, N-d dystrophic neurites and spheroids or cloudy spongiform-like alteration were supposed swelling of axon terminals showed obvious axonopathy. Compared with normal control axon in the dorsal column nucleus and spinal trigeminal nucleus, the N-d neurites dilated with aging eventually induced alteration of membrane permeability and stability because of the swelling dystrophic axons and enlarging varicosities. For diversity of formazan deposition, especially high-density core with pale halo profile, leaky of axon membrane was postulated for N-d dystrophic structure. The anomalously widespread cloudy or plaque like N-d positivity strongly remind the possibility leaky of N-d from damaged spheroid or dilated neurite. This irregular aberrant formation may still make dynamic existence. We further postulated that the N-d for neuronal dystrophy in aging rat may distribute in two locations: intracellular and extracellular localizations. Neuronal dystrophy resulted in axon membrane leaky in the medulla oblongata of aging rat, which were thought to disrupt neuronal function.

### Identical to NOS

NOS immunoreactivies is completely match with N-d staining in fish for colocalized staining[92]. NOS positive neurons and processes were seen in gracile nucleus [93, 94]. Up to date, the colocalization of NOS and N-d were not well illustrated in both young adult and aged rats. In our present study, double staining of NOS immunocytochemistry and N-d histology presented several categories: double staining neuron, single staining NOS neuron, double staining spheroid and neurite, single staining N-d spheroid and neurite. The N-d spheroid and neurite are aging-related neuronal deterioration. That the N-d histology do not completely colocalized NOS immunocytochemistry has been mentioned in some studies [21, 27–29]. NOS and N-d in rat did not show completely match as that in fish staining[92]. The colocalization of NOS immunoreactivity and N- d staining has been examined in neurons of the ileum and colon of the guinea-pig, which shows a one-to-one correlation in all neurons. However, effect of N-d activities does not change when the tissue is pharmacologically pre-treated with NOS inhibitor, N_G_-nitro-L-arginine or L-arginine or natural substrate of NOS, L-arginine[95]. However, not all of the tertiary and secondary plexuses in the stomach, duodenum and ileum show colabeling NOS-positive with N-d[96]. In the neocortex of monkey, area-specific differences are detected in the pattern of distribution of fiber plexuses[97]. We used to test the N-d dystrophic spheroid by different antibody and showed less colocalization with NOS immunocytochemistry (data not showed here). It is reasonable that N-d dystrophic spheroid and neurite may not consistently co-label with NOS. Dense or light stained spheroid and neurite indicated a different mixture of NOS and N-d bioactive components during the aging process in the sensory nuclei. The mechanism of the specific alteration is still unclear. The evidence showed that the N-d neuronal terminals was dystrophic dilation. The positivity of NOS and N-d in neuron fibers occurs more diversity than that of the neuronal soma.

### Aging dysfunction corelated with spinal cord

The dorsal column nuclei receive inputs from spinal cord[98, 99]. Besides somatic sensory input, gracile nucleus also receives ascending fiber from pelvic organs[45, 100, 101]. Aging might cause dysfunctions with voluntary function for example impaired movement and coordination especially in lower extremities and involuntary (autonomic) body functions, including urinary and bowel dysfunction such as: constipation and loss of bladder or bowel control [102], and sexual dysfunction for instance, dysfunction of erection. N-d neurodegeneration of gracile nucleus revealed a unique hallmark remarkable visceral and somatic sensory deterioration in aged rats.

Our previous study reports that the N-d fibers and IML in the lumbosacral spinal cord are vulnerable to the aging change[21, 103, 104]. However, examination of N-d histology in dorsal column and spinal trigeminal nucleus in aged pigeon fails to find massive N-d dystrophy in the gracile nucleus[105]. Compared with mammals, dominant movement for pigeon is flying. Somatic sensation and pelvic organs in pigeon are also different from that of mammals. Severe sensory-autonomic neuropathy presents accentuated central peripheral distal axonopathy and also indicate fiber degeneration in the gracile fasciculi, while the neurons in the DRG are relatively resistant to cell loss[106]. Severely dystrophic axons in animal model of Alzheimer’s disease still remain continuity and connect with cell bodies[107]. With aging, many individuals gradually suffer from muscle weakness and further gait disturbances of lower limbs. Distal sensory loss and absent ankle reflexes with age are not occurred in every aged people over 65+ year old[108]. In our research, the N-d neurodegeneration in the gracile nucleus was almost detected in every individual of aged rats. We also found that there was another aging-related dystrophic neurodegeneration such as tauopathy in the gracile nucleus and besides N-d neurodystrophy.

Axonal swellings, described in the CAJAL, BIELSCHOWSKY and others, are occurrence in the gracile nucleus[32, 33, 109] and quite diverse and manifold in their shape, size, appearance and stainability. Similarly, N-d dystrophic spheroid or neurite of our finding also showed tremendous various in their shape, size, appearance and stainability. N-d dystrophy was not only occurrence in gracile nucleus but also in cuneate nucleus as well as in spinal trigeminal nucleus. The origination of the N-d dystrophy in the gracile nucleus was partially ascending fiber of DRG neurons from lower extremities. This results strongly implicated that N-d dystrophy in the gracile nucleus was subject to aging-dysfunctional movement. We believe that increased mortality of aging was due to ground level fall because of muscle weakness and gait disturbance of lower extremities[110]. Interfering of N-d dystrophy may rescue and restore muscle weakness and gait disturbance of lower extremities which improve a fall-prevention.

The authors report no conflict of interest.

## Acknowledgments

This work was supported by grants from National Natural Science Foundation of China (81471286), Liaoning Training Programs of scientific research and career development for Undergraduates, 201410160007 and Research Start-Up Grant for New Science Faculty of Jinzhou Medical University. The authors are responsible for the content and writing of this paper.

